# Oviposition in flight: the *Sabethes albiprivus* incredible egg-throwing behavior

**DOI:** 10.1101/2020.01.31.927228

**Authors:** Genilton Vieira, Maria Ignez Lima Bersot, Glauber Rocha Pereira, Filipe Vieira Santos de Abreu, Agostinho Cardoso Nascimento-Pereira, Maycon Sebastião Alberto Santos Neves, Maria Goreti Rosa-Freitas, Monique de Albuquerque Motta, Ricardo Lourenço-de-Oliveira

## Abstract

Mosquitoes display highly variable oviposition strategies and behavior. By using a high-speed camera, we detailly documented for the first time the egg-throwing strategy of the sylvatic yellow fever vector *Sabethes albiprivus* Theobald in laboratory. An oviposition trap made with sapucaia nut for field collection of tree-hole mosquitoes and obtaining *Sa. albiprivus* eggs either in the field or in the laboratory colony is described.

## Introduction

There are more than 3,500 mosquito species (Diptera: Culicidae) described (WRBU 2019). Mosquito egg-laying strategies and behavior are highly variable (Consoli & Lourenço-de-Oliveira 1994). Females are usually inseminated very early after emergence from pupa or even during emergence, such as in genus *Deinocerites*. Thereafter, mosquito female matures eggs throughout their adult lifespans and lay egg clumps generally when completing a gonothrophic cycle following a blood meal in the case of hematophagous species, which correspond to the great majority of species (Consoli & Lourenço-de-Oliveira 1994). Gravid mosquitoes may use of a variety of water collections to lay eggs: dirty or clean; sunny or shade; easily visible or hidden, and so on according to species. At each egg-laying event, a gravid mosquito female may drop from a unique egg to a series of single eggs in few minutes or produce a batch of numerous glued eggs. Interestingly, those species that drop a single or lay isolated single eggs often disperse (skip-oviposition) between each egg-laying event scattering their progenies (Bentley & Day 1989; Consoli & Lourenço-de-Oliveira 1994). This is the egg-laying strategy of sylvatic arbovirus carrier mosquitoes of genus *Sabethes*.

Mosquitoes of genus *Sabethes* are daytime bitter insectcs that are considered for many biologists the most beautiful creatures on Earth. They exhibit an ensemble of behavioral and morphological characteristics that make them unique among the mosquitoes, like the colorful iridescent scales, especially with blue, green and violet reflexes that cover much of their bodies (Consoly & Lourenço-de-Oliveira 1994). Besides, both male and females of subgenus *Sabethes* possess an elaborate ornament mostly on mid-legs (tibia and tarsomere I) formed by differentiate elongate colorful scales that form paddle-like structures (Hancock et al. 1990a). The functions and existence of “paddles” in both sexes has been the object of several investigations on mosquito species evolution and behavior, notably in colonized *Sa*. (*Sab*.) *cyaneus* (Fabricius) (Hancock et al. 1990a,b, South et al. 2009).

Gravid females of *Sabethes* mosquitoes almost exclusively explore water collections stored in plant cavities like tree holes and pierced, cut or broken bamboo and exhibit a peculiar oviposition strategy. Instead of gently lay the egg right on the water surface or on other subtracts close the water line as the great majority of mosquitoes does, *Sabethes* species shoot the egg onto the water in flight in an extraordinary performance comprising typical fly combined with body and legs movements. This peculiar egg-laying behavior was firstly described by Galindo et al. (1955) and Galindo (1957, 1958) by observing with nude eyes and making photos of gravid females in a colony of *Sabethes chloropterus* (von Humboldt). However, the high speed of flies, body and leg movements performed by gravid females at the moment of dropping the eggs could not be ideally understood by using the limited available instruments in the 1950’s.

Life as we see it is unreal. Our limited human senses allow us only to see objects moving at a speed of more than 13 milliseconds (Potter et al 2014). And it is from this limitation that we make observations and conclusions about the world we live in. High speed capture image technologies allow the broadening of the apprehension of the reality. Reality changes, and the understanding of this reality too. In this article, using a high-speed camera, we detailly documented for the first time the unique egg-laying behavior of *Sabethes albiprivus* Theobald 1903 in laboratory, particularly at exact moment when the gravid female expel the egg and throw it to reach deep the water contained in a simulated tree hole. We also evaluated the speed of this extraordinary oviposition behavior.

*Sa. albiprivus* belongs to subgenus *Sabethes*, which comprises 18 Neotropical valid species distributed from Mexico to northern Argentina (WRBU 2019). It is a common mosquito in the Atlantic Forest from southeastern Brazil to northern Argentina, where it has been considered to play a role in yellow fever virus sylvatic transmission (Goenaga et al 2012, Couto-Lima et al 2017, Abreu et al 2019).

## Material and Methods

Mosquitoes were from a laboratory colony derived from samples of immature forms and naturally mated field-collected females from Tinguá (22° 32′ 43″ S, 43° 23′ 5″ W), Rio de Janeiro. The establishment of this colony favor previous experimental evaluation of vector competence of *Sa. albiprivus* to yellow fever virus (Couto-Lima et al 2017).

Immature *Sa. albiprivus* were collected with sets of oviposition traps consisting of fallen seedless wooden monkey pot tree (*Lecythis pisonis*) nuts partially filled with dechlorated water with the top roughly covered with the wooden fruit lid to leave a small opening (Figure 1). To slow the loss of water contained in the sapucaia nut, we applied a layer of ethylene-vinyl acetate (E.V.A.) hot melt adhesive to the basal outer surface. *L*. *pisonis* is a nut producing tree native to the Amazon very common to urban parks and forest in Brazil, also called cream nut, paradise nut or sapucaia nut. Traps were hung in the forest canopy for two weeks. Then, obtained immature mosquitoes were screened to species and reared until adult stage in the laboratory as previously described (Couto-Lima et al 2017). The *Sa. albiprivus* colony used in this study has been maintained at our laboratory for several generations since 2013, at 26 ± 1°C, 70–80% RH and a 12L:12D photoperiod, at a population size of around 100 adult individuals per cage. Mosquito collection and colonization were respectively approved by local environmental authorities (SISBIO-MMA licenses 37362-2) and the Institutional Ethics Committee on Animal Use (CEUA-IOC license LW-34/14) at the Instituto Oswaldo.

**Figure 1:**
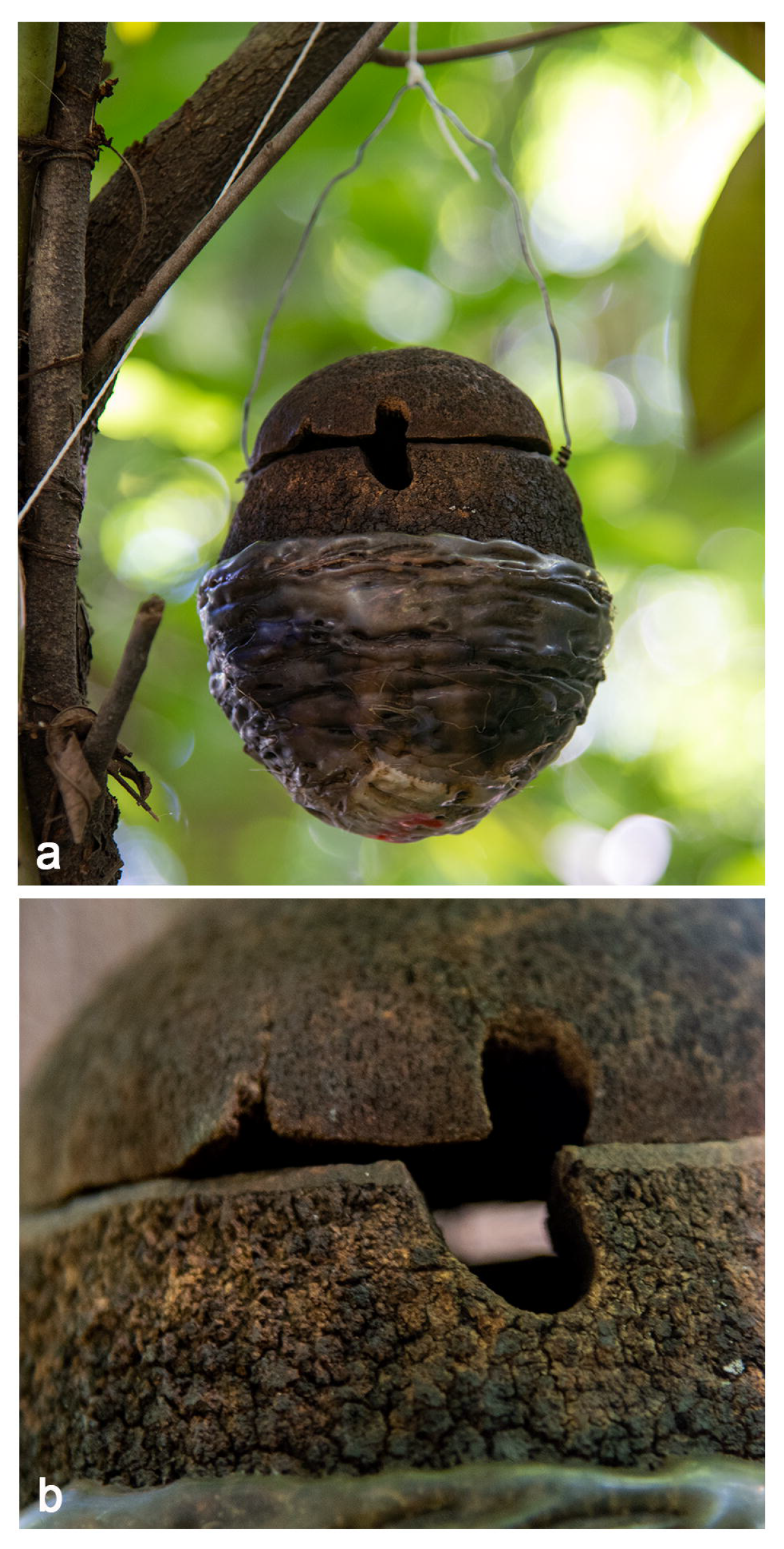
The sapucaia nut oviposition trap. (a) The trap hung at the forest canopy, with the two-thirds of basal outer nut surface covered with a layer of ethylene-vinyl acetate E.V.A. hot melt adhesive to slow the loss of water contained. (b) A detail of the uneven closure made by the nut lid, leaving a narrow outward opening through which gravid *Sabethes* mosquitoes throw their eggs, both on the field and at the laboratory.

In this study, the sapucaia nut oviposition trap was also used in the laboratory to simulate a deep tree hole, the natural larval habitat of *Sa. albiprivus* and other related *Sabethes* species (Galindo et al 1955) to allow for observations.

The egg-laying behavior of *Sa. albiprivus* was shoot with a Phantom® Miro 310 one-megapixel camera capable of recording until 3,200 frames-per-second at full 1280 × 800 resolution. We also have worked with especial lenses capable of shooting and taking photographies at relatively small distance with great resolution. The shootings were made in a glass cage where sets of *Sa. albiprivus* gravid females had been released 1-2 days previously.

## Results

The extraordinary egg-laying behavior of *Sa. albiprivus* is illustrated with video sections and a sequence of eight selected frames from one movie displayed in figures. The oviposition steps are described below.

### Seeking to identify the entrance of the tunnel through which the egg will be launched towards the water

Females lay eggs one at a time (Figs. S1-3). Gravid females approach the opening and performs a sequence of rapid and short up and down flights. Notice that the egg to be launched is already hold at the abdomen tip. The egg of *Sa. albiprivus* and other *Sabethes* is not dropped immediately after leaving from the ovipositing tube and vagina (gonotreme). Contrarily, after extrusion, it remains hold at the abdomen tip assisted mainly by the cerci and postgenital lobe until being thrown. At this time, all three pairs of legs essentially display the characteristic positions of a flying *Sabethes* mosquito (hovering, like gliding in the air) (Figs. S3 and 2a): former legs prominently directed upward above the thorax, perpendicular to the body main axis, with tarsomeres curved backward; mid-legs directed downward and outward, forming an angle of 70-80° between each other; hid-legs directed to the rear with tarsomeres visibly curved frontward. Body axis (proboscis, thorax and abdomen) forming an angle of around 110° (Fig. 2a).

**Figure 2:**
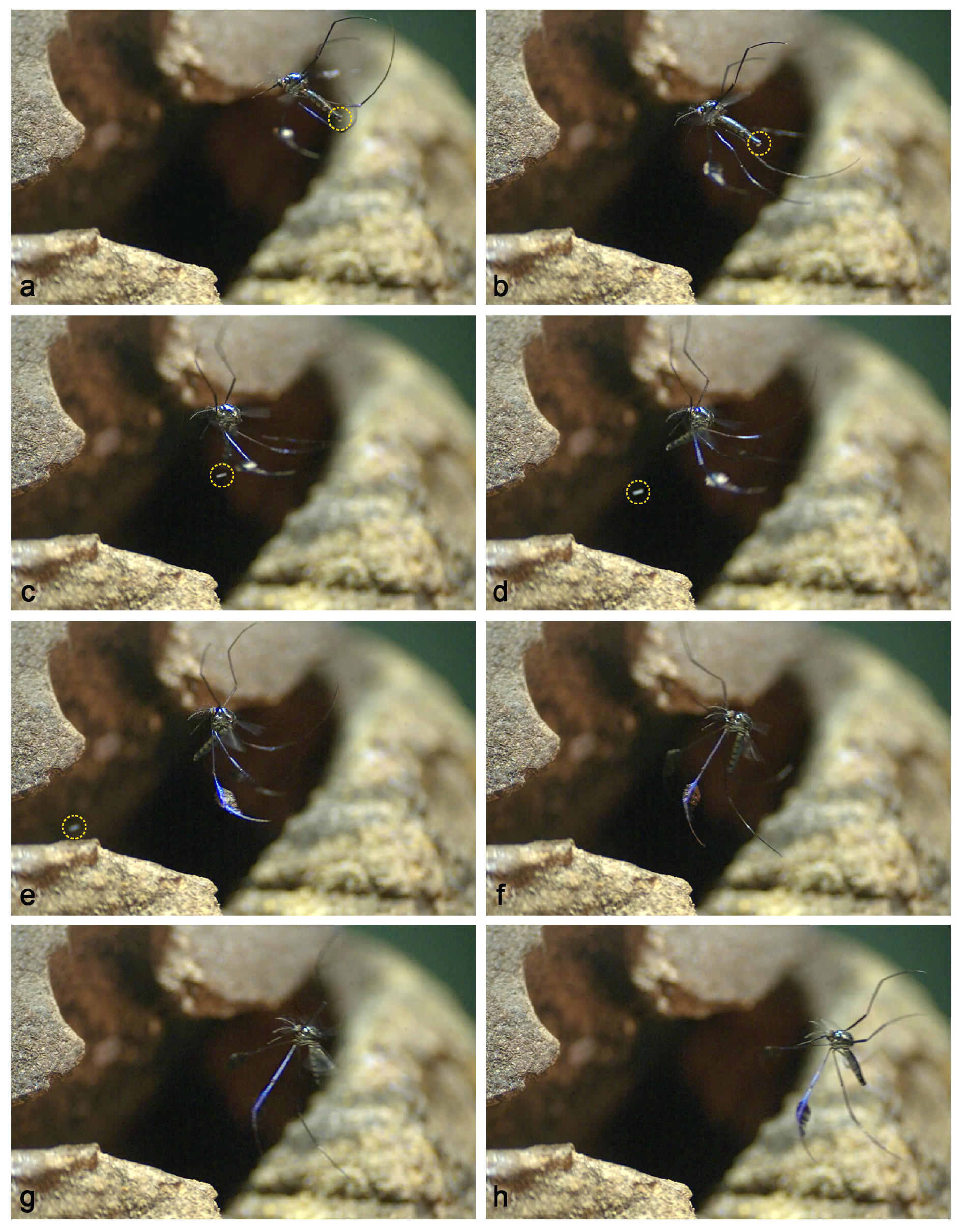
Sequence of frames taken from a *Sabethes albiprivus* gravid females laying an egg in a sapucaia nut trap in the laboratory filmed at 3200 frames per seconds. They illustrate the distinct body and leg movements made before, during and after the egg-throwing. Circles underline the egg still hold at the abdomen tip (a,b) and traveling toward the hole (c-e).

### Getting ready to throw the egg

Female break the up and down flight, and hovering the opening, begins to make movements with legs before shooting the egg. Hind-legs become less curved and move downward and frontward (Fig. 2b). Then, the female is ready to shoot the eggs. It almost simultaneously twitches the head downward and backward and move the mid-leg backward and fore-leg frontward (Fig. 2c); fore-legs may touch the wood framing the opening through which the egg will be launched towards the water (Fig. S3). At this moment, female strongly start thrusting the whole abdomen forward (Figs. S3 and 2c).

### Throwing the egg

The egg is vigorously thrown (Figs. S2 and S3). The mosquito body fold downward and body main axis assumes a small angle ventrally; the proboscis tip almost touches the abdomen (Fig. 2d). The egg speedily travels through the opening and goes deep into the hole, while the female remains outside the opening (Figs. S1-4 and 2e). We calculated the egg travel speed when filming some *Sa albiprivus* females with a Phantom camera at a speed of 1500 frames per second (or 0.04 s per frame). Tanking the 3mm abdomen length as calibration and measuring the distance traveled by the eggs until the moment when they disappear in the hole (12.0-13.9), we found that the egg travel speed is almost 1 m/s (0.934-0.998 m/s). Female may fail to throw the egg into the hole. Rarely, the egg may not be released at the time of throwing and fall from abdomen tip just after (Fig. S4), or even falling from the abdomen tip before attempting to throw (Fig. S5). In these cases, the egg may fall out and be lost or may eventually bounce and roll into the hole (Figs. S4 and S5).

### Flying back and getting ready to lay another egg

The mission achieved, female prepares to fly back. The fore- and mid-legs quickly and prominently move forward and body starts unbending (Fig. 2f). Female rapidly flies back. The rear flight seems to gain force when it twists the mid-leg axis so that paddle faces forward while switching fore- and hid-legs position to backward (Figs. S2-4 and 2g). Female’s body and legs almost resume the typical shape and position and restarts typical hovering *Sabethes* flight (Figs. S1, S3 and 2h). It can be noticed that another egg rapidly is extruded from vagina and is available at the abdomen tip to be thrown (Fig. S1).

## Discussion

Insofar as we know it is the first time the egg-laying strategy and behavior and the contribution of corporal and leg movements are recorded in such details for a *Sabethes* mosquito. Also noteworthy is the originality of the description of the use of an oviposition trap made with sapucaia nut for field collection and egg-laying of *Sa. albiprivus* and other tree-hole mosquitoes.

The sapucaia nut oviposition trap had great success as a tree hole simulator, both in the laboratory and on the field. In the laboratory, gravid *Sa. albiprivus* females readily searched for the small opening provided by the barely fitted fruit lid as shown in the videos. It agrees with the previous reports of the propensity of *Sabethes* to search for tree-holes with small and laterally oriented cryptic entrance (Mattingly 1969, Mangudo et al. 2014), and refusing those with water surfaces directly exposed (Yanoviak 1999). Traditional ovitraps made with black plastic containers are not useful for obtaining *Sabethes* eggs either in the field or in the laboratory (Yanoviak 2001). In the field, the sapucaia nut trap we developed successfully collected other mosquito species that oviposit tree holes with a narrow external opening besides *Sa. albiprivus*, such as *Sa. purpureus* (Theobald), *Sa. belisarioi* Neiva, *Sa. batesi* Lane & Cerqueira, *Sa. chloropterus, Sa. gynothorax* Harbach & Petersen, *Sa. tarsopus* Dyar & Knab, *Sa. glaucodaemon* (Dyar & Shannon) and *Wyeomyia nigritubus* Galindo, Carpenter & Trapido. However, when the lid was accidentally dislocated or dropped on the forest ground during the trap operation, leaving a large opening to the outside or fully exposing the water contained in the sapucaia nut, a clear qualitative change in mosquito fauna composition is noticed: instead of aforementioned species, other tree-hole mosquitoes are collected, such as *Limatus pseudomethysticus* (Bonne-Wepster and Bonne), *Haemagogus janthinomys* Dyar, *Haemagogus leucocelaenus*, (Dyar & Shannon), *Culex (Carrollia)* sp. and *Aedes terrens* (Walker).

Concerning oviposition behavior and strategy in *Sabethes*, in the 1950’s, Galindo and colleagues (Galindo et al. 1955; Galindo 1957, 1958) have explored and photographed *Sa. chloropterus* throwing eggs through holes drilled in bamboos, but most details of *Sabethes* egg-laying could not be evidenced. For instance, with the slow-motion video we could determine the egg thrown by a gravid *Sa. albiprivus* travels in a speed of almost 1 m/s. It also evidenced that legs, especially the mid-legs ornated with the paddles play an important role in the egg-throwing strategy of *Sa. albiprivus* and may of other species of *Sabethes*. Hancock et al. (1990a) concluded that paddle removal in *Sa. cyaneus* does not change oviposition behavior or flight and proposed that the paddles would not help in hovering or rapid reverse flight. For these authors (Hancock et al. 1990b) paddles are implicated in female mate attraction as insemination rates are significantly lower in paddleless *Sa. cyaneus* females. Although, functions of paddles remain undetermined, we noticed that at least the reverse flight after egg-throwing seems to gain force at the time *Sa. cyaneus* females twists the mid-leg axis so that paddle faces. Although the egg of *Sa. albiprivus* can travel in a substantial speed, the gravid female throws it very close the entrance of the tunnel leading the water containing cavity, with its fore-legs sometimes touching the external opening. This behavior has also been described for *Sa. chloropterus* whose females hover in front of the opening at a distance of a few millimeters to as much as 5cm before eject the egg (Galindo 1958). Maybe the force and speed with which the egg is thrown by *Sabethes* mosquitoes are not enough to ensure the to travel long distances from the point of ejection. In fact, Galindo (1957) suggested that the egg of *Sa. chloropterus* travel from 2.5 to 10 cm in a straight. In any case, the peculiar rhomboid shape of many Sabethini’s eggs (Galindo 1958, Barr & Barr 1969) may facilitate their reaching the water, by rolling and falling under gravity. Because of this egg morphology and the action of gravity, it is possible that even the eggs whose throwing was flawed as we have recorded in video for *Sa. albiprivus* may eventually reach the water deep in the hole and ensure hatching it dropped into the opening. Even son, natural selection may favor females who throw better, maximizing the hatching chances of their eggs.

In mosquitoes, the oocyte half emerges through the gonotreme and lasts for few seconds held by the lateral egg guides and postgenital lobe to be fecundated (Clements 1999). Then, the egg is immediately dropped in the great majority of mosquito species. However, as we evidenced for *Sa. albiprivus*, following fecundation of the partially extruded oocyte, the egg is hold at the abdomen tip until the gravid female decides to throw it through the hole. That is, after passing the gonotreme, the egg of *Sa. albiprivus* is not immediately dropped, thrown or flicked, a behavior probably shared with other Sabethini and insect to other orders like stick insects (Phasmatodea) and dragon-flies (Odonata) (Carlberg 1983). Also, we observed that *Sa. albiprivus* females lay eggs one at a time and are ready to throw another egg a few seconds afterwards. But, according to Galindo (1958) *Sa. chloropterus* would shoot one or two eggs through the hole drilled in bamboo.

Therefore, thank to high speed image capture technologies we could uncover the amazing details of the egg-laying behavior of *Sabethes* mosquitoes, a neotropical genus involved in the transmission of yellow fever virus. In addition, we develop an oviposition trap to almost selectively collect these mosquitoes.

## Authors’ contributions

MAM, GV, MILB and RLO conceived and designed the research, GV filmed and prepared the videos and figures, FVSA, MSASN, MAM, ACNP, RLO and GRP collected mosquitoes to initiate the colony and tested the sapucaia-nut oviposition trap, MILB and MAM colonized mosquitoes in laboratory, RLO and MGRF analyzed images and wrote the manuscript. All authors reviewed and approved the manuscript

## Acknowledgements

To Marcelo Q. Gomes for the support in the field, and Heloisa Diniz and Alexandrp M. Freitas for the help with illustrations.

## Legends of figures

Figure S1: *Sabethes albiprivus* gravid females laying eggs in a sapucaia nut trap in the laboratory filmed at normal speed (~30 frames per second). Notice the sequence of rapid and short up and down flights, and the egg to be thrown hold at their abdomen tip.

Figure S2: The egg-throwing behavior of gravid *Sabethes albiprivus* mosquitoes in laboratory filmed at 3200 and 1500 frames per second. Notice the movements of distinct pairs of legs before and during egg-throwing into the hole. The rear flight seems to gain force when the female twists the mid-leg axis so that paddle faces forward while switching fore- and hid-legs position to backward.

Figure S3: *-* The egg-throwing behavior of gravid *Sabethes albiprivus* mosquitoes in laboratory filmed at 2400 frames per second. Notice the up and down flight and hovering executed by female very close to the opening. Forelegs touch the opening of the sapucaia trap when the egg is ejected.

Figure S4: *-* A gravid *Sabethes albiprivus* mosquitoes may fail in throwing the egg. The egg falls after the attempt; a new egg is soon extruded and is noticed at the abdomen tip. Filmed at 1000 frames per second.

Figure S5: The egg may fall before the attempt of throwing it by a gravid *Sabethes albiprivus* mosquito. The egg may eventually fall into the hole. Filmed at 3200 frames per second.

